# Naked Mole-Rat Cells are Susceptible to Malignant Transformation by *SV40LT* and Oncogenic Ras

**DOI:** 10.1101/404574

**Authors:** Fazal Hadi, Yavuz Kulaberoglu, Kyren A. Lazarus, Paul Beattie, Ewan St. John Smith, Walid T. Khaled

## Abstract

The Naked Mole-Rat, *Hetercephalus glaber*, is a mouse-sized subterranean rodent native to East Africa. Research on NMRs is intensifying in an effort to gain leverage from their unusual physiology, long-life span and cancer resistance. Few studies have attempted to explain the reasons behind NMRs cancer resistance, but most prominently Tian et al. reported that NMR cells produce high-molecular weight hyaluronan as a potential cause for the NMR’s cancer resistance. Tian et al. have shown that NMR cells are resistant to transformation by SV40 Large T Antigen (*SV40LT*) and oncogenic HRAS (*HRAS*^*G12V*^), a combination of oncogenes sufficient to transform mouse and rat fibroblasts. We have developed a single lentiviral vector to deliver both these oncogenes and generated multiple cell lines from five different tissues and nine different NMRs, and report here that contrary to Tian et al.’s observation, NMR cells are susceptible to oncogenic transformation by *SV40LT* and *HRAS*. Our data thus point to a non-cell autonomous mechanism underlying the remarkable cancer resistance of NMRs. Identifying these non-cell autonomous mechanisms could have implications on our understanding of human cancer development.

---

The Naked Mole-Rat (NMR), *Hetercephalus glaber*, is a mouse-sized subterranean rodent native to East Africa. Research on NMRs is intensifying in an effort to gain leverage from their unusual physiology^1^,^2^, long-life span^3^ and cancer resistance^2^. NMR’s acclamation as a cancer resistance species mainly derives from studies describing a lack of spontaneous neoplasms in captive NMR colonies despite the great number (>2000) of reported necropsies^4^. Although two recent studies described a very small number of cases of spontaneous tumours in captive NMRs^5,6^, the NMR’s cancer resistance is still remarkable relative to their size and life-span. Few studies have attempted to explain the reasons behind NMRs cancer resistance^7–9^ and most prominently Tian et al. reported that NMR cells produce high-molecular weight hyaluronan as a potential cause for the NMR’s cancer resistance^10^. With the publication of the NMR genome^11,12^ and advances in the CRISPR/Cas9 gene editing technology^13^ we set out to identify NMR genes responsible for their cancer resistance. We based our approach on Tian et al.’s observation that NMR cells are resistant to transformation by SV40 Large T Antigen (*SV40LT*) and oncogenic *HRAS* (*HRAS*^*G12V*^), a combination of oncogenes sufficient to transform mouse and rat fibroblasts^14,15^.

We therefore, generated a number of lentiviral vectors which would allow us to deliver sgRNAs along with *SV40LT* and/or *HRAS*^*G12V*^ under the control of two different promoters (*PGK and EF1α*, Figure 1a). As reported by Tian et al., we expected that NMR cells transduced with any of our vectors would not grow in anchorage-independent conditions unless a further gene was perturbed, thereby providing us with a defined system for our studies. Using our lentiviral vectors, we generated over 90 cells lines from 5 different tissues (intestine, kidney, pancreas, lung and skin) from 9 different NMRs (Figure 1b and Supplementary Table 1). We then proceeded to test few cell lines for anchorage independent growth as described by Tian et al. As expected, and in line with previous reports, primary cell lines and those transduced with *SV40LT* alone failed to grow in soft agar even after six weeks in culture whereas, to our surprise and contrary to previous reports, NMR cell lines expressing both *SV40LT* and *HRAS*^*G12V*^ formed robust colonies in soft agar (Figure 1c-d and Supplementary Figure 1). These results were reproducible for all cell lines tested, irrespective of the animal or promoter used to drive the expression of the exogenous *SV40LT* and *HRAS*^*G12V*^ genes (Figure 1c-d and Supplementary Figure 1).

**Figure 1:**
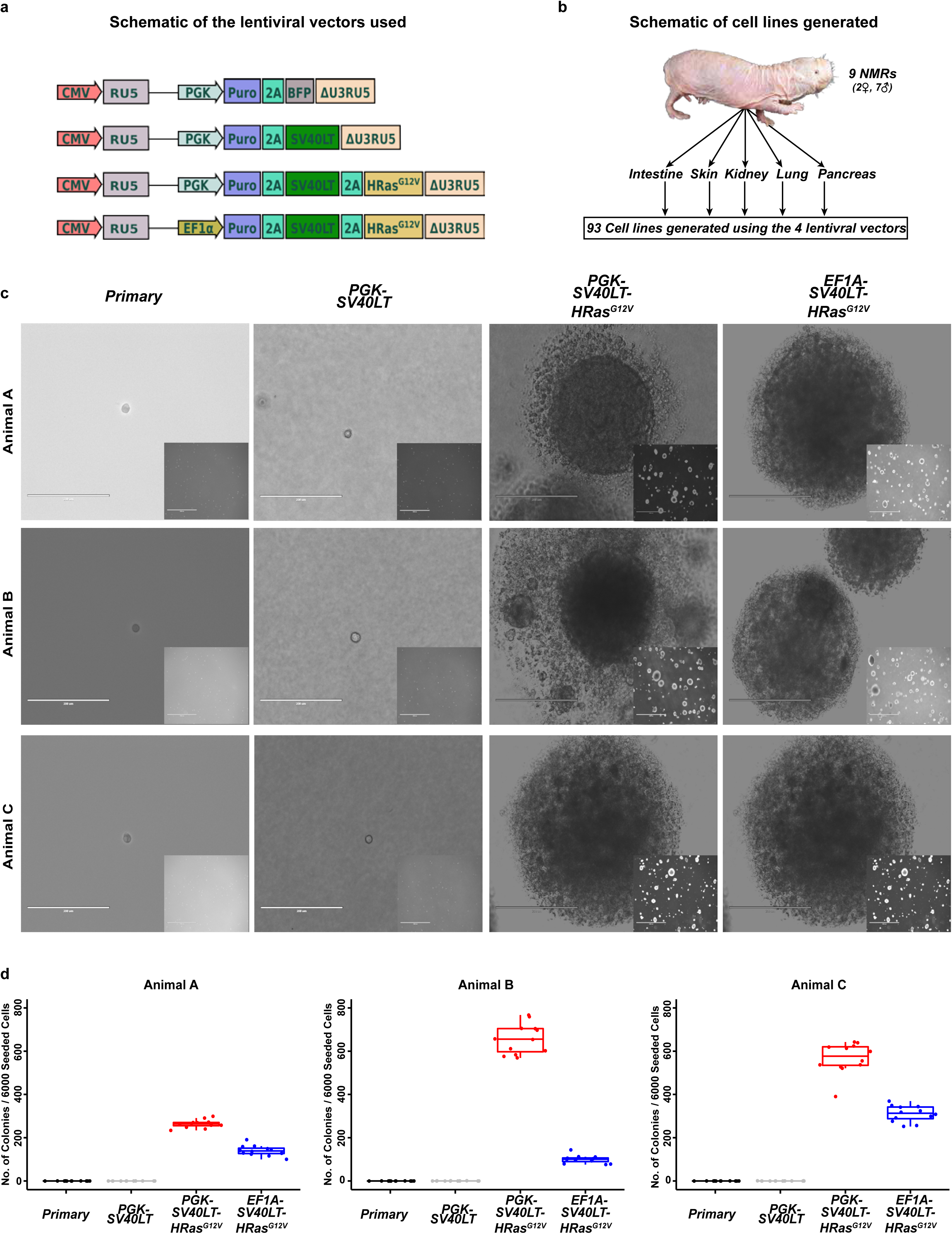
**(a)** Schematic of the lentiviral vectors generated and used in this study. **(b)** Schematic representation of the different cell lines generated as part of this study. **(c)** Representative images of the colonies observed in the soft agar assay. Each row represents the skin cell lines from one animal and each column represents the cell lines transduced with one of the four vectors described in **(a)**. Inset images are a lower magnification (scale bar = 2000µm) of the same field of the larger image (Scale bar = 400µm). **(d)** Quantification of the colony numbers from the soft agar assay in **(c)**. The soft agar assay for each cell line was repeated twice and on each occasion six technical replicates were performed. Each dot represents the number of colonies observed from an individual technical replicate. In total an excess of 2500 images were quantified using ImageJ.

Based on these results, we next decided to test the tumourigenic potential of the cell lines expressing *SV40LT* and *HRAS*^*G12V*^. Non-transduced parental cell lines and their respective *SV40LT* and *HRAS*^*G12V*^ overexpressing cell lines were injected subcutaneously into NSG immunocompromised mice and monitored for tumour growth. As early as day 4 post-injection, tumour masses were detected in animals bearing cell lines expressing *SV40LT* and *HRASG*^G12V^ (Figure 2a, b and Supplementary Figure 2a). By day 39 post-injection all animals bearing transformed lines had been collected (Figure 2b), whereas animals injected with non-transduced cells did not show any signs of tumour growth even 60 days post-injection when the experiment was terminated (Figure 2a).

**Figure 2:**
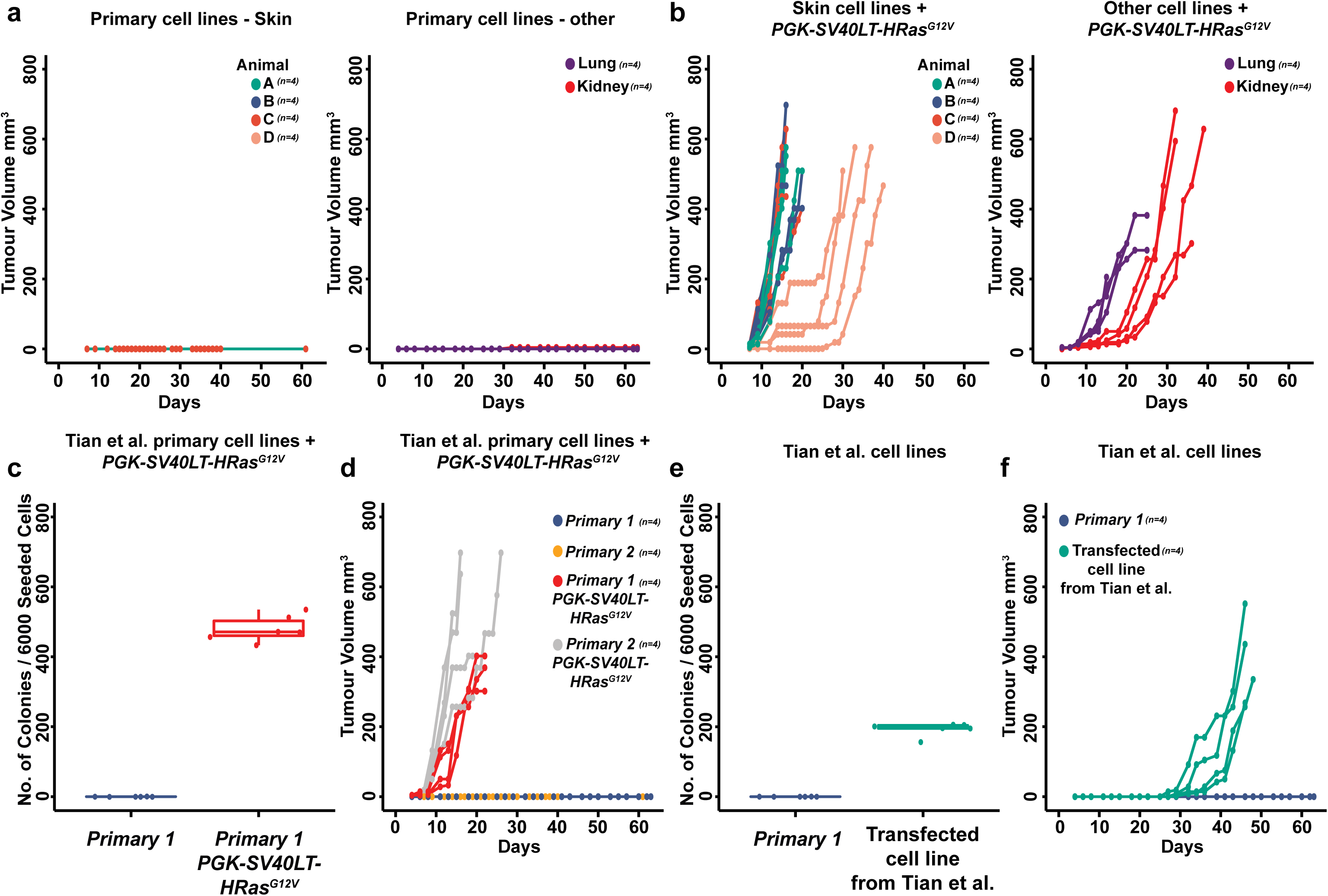
Quantification of xenograft tumour growth from NSG mice injected with **(a)** primary cell lines or **(b)** their respective *PGK-SV40LT-HRAS^G12V^* transduced cell lines. Each cell line was injected into four mice each, each mouse being represented by a single line on the graph. **(c)** Quantification of the colony numbers observed from a primary skin cell line obtained from Tian et al. and its respective *PGK-SV40LT-HRAS^G12V^* transduced cell line. Each dot represents the number of colonies observed from an individual technical replicate. **(d)** Quantification of xenograft tumour growth from mice injected with two primary skin cell lines obtained from Tian et al. and their respective *PGK-SV40LT-HRAS^G12V^* transduced cell lines. **(e)** Quantification of the colony numbers observed from a primary skin cell line obtained from Tian et al. and its respective (*SV40LT and HRAS^G12V^*) transfected cell line generated by Tian et al. Each dot represents the number of colonies observed from an individual technical replicate. **(f)** Quantification of xenograft tumour growth from mice injected with the two cell lines described from **(e)**.

In an attempt to explain the discrepancy between our findings and those of Tian et al. the authors provided us with detailed list of materials and protocols which we used throughout our study. In addition, the authors also provided us with primary and transformed cell lines to test in our lab. Similar to our primary cells from multiple tissues, primary skin cell lines obtained from Tian et al. were transformed by our *SV40LT* and *HRAS*^*G12V*^ overexpressing vectors and formed robust colonies *in vitro* and tumours in xenograft assays (Figure 2c, d and Supplementary Figure 2b-d) thus, excluding any variation in NMR colonies as the cause for the differences observed. Interestingly, cells transformed by Tian et al. also formed colonies *in vitro* and tumours in xenograft assays which we have quantified (Figure 2e, f and Supplementary Figure 2b-d). However, these cells had a lower clonogenic efficiency and longer tumour latency.

In our opinion, the main reason behind the discrepancy in the data is that Tian et al. used multiple transient expression vectors to transfect the primary NMR cells without robust antibiotic selection. Given the transformative nature of the exogenous oncogenes it is inevitable that some cells will become transformed and be selected for over-time, but at a much lower rate thus, explaining the lower colony efficiency and tumour latency we observed with their cell lines. In contrast, we used a single lentiviral vector to deliver both *SV40LT* and *HRAS*^*G12V*^ into the cells and generated stable cell lines. Our vectors carry a puromycin resistance gene, which was used to select for stable cell lines from the outset, thus explaining the robust nature of the colony assays and tumour growth.

It is important to note that we do not question the finding by Tian et al. regarding the presence of high molecular weight hyaluronan in NMR cells, in fact we have also detected this in our cells. However, the importance of high molecular weight hyaluronan in NMR cancer resistance needs to be reassessed in the light of our findings. In addition, one of the transformation vectors (*HRAS*^*G12V*^) and the *Has2* shRNA vector used by Tian et al. carry the same antibiotic resistance gene making it difficult to interpret the results.

Finally, based on our results we conclude that NMR cells are not resistant to transformation by *SV40LT* and *HRAS*^*G12V*^. Our data thus suggest that the key mechanisms behind NMR cancer resistance are non-cell autonomous and instead might be explained by a unique microenvironment and/or immune system. It is important that the field is aware of our findings so that efforts in understanding the NMR’s remarkable biology are channelled in the right direction.

## Materials and methods

### Animals

All experimental animal work was performed in accordance to the UK Animals (Scientific Procedures) Act 1986 Amendment Regulations 2012 and approved by the Animal Welfare and Ethical Review Bodies at the University of Cambridge and the Wellcome Trust Sanger Institute. Various tissues (see below) from adult male and female NMRs aged between 37 and 244 weeks were used in this study (see Supplementary Table 1 for weight, sex and age information of each NMR used).

### NMR Primary Cell Isolation

Following CO_2_ exposure and decapitation, five different tissues namely skin, pancreas, lung, kidney and intestine were collected from each animal on ice-cold NMR Cell Isolation Medium (DMEM high glucose (Gibco # 11965092) supplemented with 100 units ml^-1^ Penicillin, 100µg ml^-1^ Streptomycin (Gibco # 15140122)). Skin came from either underarm or underbelly area and was cleared of any fat or muscle tissue and generously sprayed with 70% ethanol. Intestines were collected from the jejunum to the ileum and cleared of any faeces by flushing with phosphate-buffered saline (PBS) before removing any fat tissue. Similarly, fat tissue and adrenal glands were removed from kidneys. Once cleared each tissue was washed twice with PBS before finely mincing with sterile scalpels. The minced lung tissue was then washed with 4ml NH_4_Cl solution (containing 0.8% NH_4_Cl in H_2_O (w/v) and 1mM EDTA, buffered with KHCO_3_, final pH7.2 - 7.6) to lyse any red blood cells. Minced skin, pancreas, kidney, intestine and lungs cleared of blood cells were then mixed with 5ml of NMR Cell Isolation Medium containing 500µl of Cell Dissociation Enzyme Mix (10mg ml^-1^ Collagenase (Roche # 11088793001), 1000Units ml^-1^ Hyaluronidase (Sigma # H3506) in DMEM high glucose (Gibco # 11965092)) and incubated at 37°C for 3 – 5 hours. Each tissue was briefly vortexed every 30 minutes to aid cell dissociation and inspected for cell dissociation. In order to reduce cell death, intestinal tissue was incubated with NMR Cell Isolation Medium containing Cell Dissociation Enzyme Mix for only 1 hour and then vigorously pipetted to dissociate cells. After complete dissociation, cells were pelleted by centrifuging at 500 g for 5 minutes and resuspended in NMR Cell Culture Medium (DMEM high glucose (Gibco # 11965092) supplemented with 15% fetal bovine serum (Gibco), non-essential amino acids (Gibco # 11140050), 1mM sodium pyruvate (Gibco # 11360039), 100units ml^-1^ Penicillin, 100µg ml^-1^ Streptomycin (Gibco # 15140122) and 100µg ml^−1^ Primocin (InvivoGen # ant-pm-2)). This cell suspension was passed through 70µm filter (Corning # 352350) and seeded on treated cell culture dishes (Greiner Bio-One). Each cell suspension was seeded on 1x 75 cm^2^ dish and 7x 25 cm^2^ dishes and incubated in a humidified 32°C incubator with 5% CO_2_ and 3% O_2_. The cells cultures on 25 cm^2^ dishes were used for cell line generation (see below) whereas those on the 75 cm^2^ dish were expanded and frozen as primary cells within four passages.

### Lentiviral Plasmid Construction

*pKLV2-U6gRNA5(BbsI)-PGKpuro2ABFP-W* (Addgene # 67974)^16^ was used as backbone plasmid for generating the *PGK-SV40LT-HRAS^G12V^* immortalisation vector. DNA fragments coding for the *T2A-SV40LT-T2A-HRAS^G12V^* sequence was synthesized using GeneArt Gene Synthesis service (ThermoFisher Scientific). The synthesized DNA fragment included the sequence encoding for the last 9 amino acids of the puromycin resistance gene and SexAI restriction site at the 5’ end and the NotI restriction site at the 3’ end to allow for direct cloning into *pKLV2-U6gRNA5(BbsI)-PGKpuro2ABFP-W*. Since the SexAI restriction site is methylation sensitive, Dam and Dcm deficient *E*. *coli* (NEB # C2952I) were used for cloning. In order to generate the PGK-*SV40LT* only vector, the T2A-*SV40LT* sequence was PCR amplified using a primer pair carrying the SexAI and STOP-NotI restriction site. The PCR fragment was digested with SexAI and NotI enzymes and ligated to the SexAI and NotI digested backbone plasmid (Addgene # 67974). The *EF1A-SV40LT-HRAS^G12V^* vector was constructed by replacing the *PGK* promoter with human *EF1A* promoter using Gibson assembly (NEBuilder HiFi DNA Assembly Cloning Kit (NEB # E5520S)).

### Cell Culture

All NMR cells (both primary and immortalised) were cultured in NMR Cell Culture Medium (as described above) in a humidified 32°C incubator with 5% CO_2_ and 3% O_2_ (Binder CB160). Media was changed twice a week and upon confluency cells were washed with PBS and detached using 1XTrypsin-PBS (Trypsin-EDTA (Sigma # T4174-100ml). After detaching cells, trypsin was inactivated by addition of NMR Cell Culture Medium and centrifuged at 450 g for 5 minutes to remove the trypsin containing medium. The cell pellet was resuspended in NMR Culture Medium and seeded on appropriate size flasks. HEK-293FT cells were cultured in DMEM high glucose (Gibco # 11965092) supplemented with 10% fetal bovine serum (Gibco) and 100units ml^-1^ Penicillin, 100µg ml^-1^ Streptomycin (Gibco # 15140122) in a humidified 37°C incubator with 5% CO_2_.

### Lentivirus Production

HEK-293FT cells were used as packaging cells for producing lentiviral particles. One day before transfection, HEK-293FT cells were seeded on 10 cm dishes (Greiner # 664960) coated with 0.1% gelatin (Sigma # 1393-100ml) in PBS (w/v) for 30 minutes. The next day, cells were cotransfected with one of the immortalization (or control) vectors (described above) and second-generation lentivirus packaging plasmids: pMD2.G (Addgene # 12259), psPAX2 (Addgene # 12260) as follows. For each 10 cm dish, 12µl of Plus reagent (Gibco # 15338100) was mixed with 5.4µg immortalization (or control) vector, 5.4µg psPAX2 and 1.2µg pMD2.G in 3ml of Opti-MEM I (Gibco # 31985062). This mixture was briefly vortexed and then incubated at room temperature for 5 minutes before adding 36µl Lipofectamine LTX (Gibco # 15338100), followed by vortexing and incubating for further 30 minutes at room temperature. Just before adding the transfection mixture, fresh 5ml (DMEM high glucose (Gibco # 11965092) supplemented with 10% fetal bovine serum) was added to the cells. Next, the transfection mixture was added to the cells and incubated at 37°C in a humidified 5% CO_2_ incubator. After 5 hours, the medium containing the transfection mixture was removed and replaced with 10ml fresh media (DMEM high glucose (Gibco # 11965092) supplemented with 10% fetal bovine serum). The cells were then incubated in a 37°C humidified 5% CO_2_ incubator for 36 – 48 hours. Following the incubation, the lentiviral particle-containing medium was collected and passed through a 0.45µm Mixed Cellulose Ester (MCE) filter (Millipore # SLHA033SS) to remove any cellular debris. Three volumes of this cleared lentiviral particle-containing medium was combined with one volume of Lenti-X Concentrator (Clontech # 631232) and mixed gently before incubating overnight at 4°C. The Lenti-X Concentrator – lentiviral mixture was centrifuged at 1500 g for 45 minutes at 4°C. The supernatant was discarded and the viral pellet was resuspended in a ratio of 1:25 of the original volume in antibiotic free media and 200 µl aliquots were stored at –80°C.

### Lentiviral Transduction

Primary NMR cells were seeded on 25 cm^2^ dishes and when the cells reach a minimum of 20% confluency, the cells were washed with PBS and 5ml of fresh NMR cell culture media supplemented with 200-250µl of defrosted lentiviral particles were added. 72 hours post-transduction, the cells were washed with PBS and fresh NMR cell culture media was added. The cells were allowed to recover for another 24 hours before starting selection with puromycin (Sigma # P7255-100MG) at a final concentration of 2µg ml^-1^ for 4 days for NMR skin, pancreas, lung and intestinal cells. For the kidney cell lines 4µg ml^-1^ of puromycin was used for 3 days. In each case, a flask of non-transduced cells was used as a control to ensure for antibiotic selection.

### Anchorage-Independent Growth (Soft Agar Assay)

NMR cell lines generated using the lentiviral vectors (above) and their primary counterparts were tested for anchorage-independent growth by seeding them in soft agar. One day before the assay was performed, 0.7g and 1g of Difco Noble Agar (BD Bioscience # 214220) was added to 100ml of deionised water to make 0.7% and 1% agar suspensions. These were autoclaved in a benchtop autoclave (Classic Prestige Medical) before storing at 4°C. The 1% agar was melted in a microwave and incubated at 37°C. Melted 1% agar and Soft Agar Medium (2X MEM (Gibco # 11935046) supplemented with 15% fetal bovine serum (Gibco), non-essential amino acid (Gibco # 11140050), 1mM sodium pyruvate (Gibco # 11360039) and 100units ml^-1^ Penicillin, 100µg ml^-1^ Streptomycin (Gibco # 15140122)) were mixed in equal volumes and 1.5ml of which was added into each well of a 6-well plate to make a bottom layer of 0.5% agar. This layer was allowed to solidify at room temperature in a cell culture hood for 2 hours. Once solidified, NMR cells were trypsinised, counted and diluted in Soft Agar Medium. The cells were then mixed in a 1:1 ratio with 37°C, 0.7% agar making a single cell suspension of 4000 cells per ml. 1.5ml of the cell-agar suspension (or 6000 cells) was added on top of the solidified bottom layer prepared earlier and allowed to solidify at room temperature for 2 hours before being supplemented with 1ml of NMR media and incubated in a humidified 32°C incubator with 5 % CO_2_ and 3 % O_2_ and 6 wells were seeded per condition per line. Cells were incubated for 6 weeks to permit colony formation and the medium on top was changed twice a week to replenish nutrients. To quantify the number of colonies observed, 20 images per well were captured in a single focal plane using a 1.25x objective on a EVOS FL2 (Thermo Scientific) thus, imaging the majority of the well. This amounted to a collection of approximately 2500 images in total which were analysed and quantified using an ImageJ macro. A random selection of images was counted manually to verify the results obtained by the macro.

### Xenograft Assay

Trypsinized cells were counted and 6 million cells were resuspended in 450 µl of HF (HBBS (Gibco # 24020091) + 1% fetal bovine serum (Gibco)). The cell suspension was then mixed with 150 µl ice-cold matrigel (Corning # 354230) before injection to get a final cell suspension of 6 million cells in 600 HF - 25% matrigel. 100µl of the cell suspension (or 1 million cells) were injected subcutaneously at the flank of NSG mice using a 25-gauge needle. Animals were monitored for tumour growth and tumour volume was measured every 48-72 hours using calipers. Cells were allowed to grow in the animals for a maximum 9 weeks (63 days) post injection or till the tumour reached a maximum allowed area of 1.2 cm^2^. Importantly the experiment was double blind - cells were assigned random names by one experimenter who handed them over to a second experimenter who then injected the animals and tumours were measured by an animal technician who was unaware of the identity of the lines.

### Genotyping PCR

NMR cells were checked for contamination by human or mouse cells using genotyping PCR. The following loci were used: *Rosa26* (mouse specific), *IL32* (human specific) and *Eras* (NMR specific). Cells were detached and pelleted as described above and genomic DNA was extracted using QIAamp DNA Blood Mini Kit (Qiagen # 51104). PCR amplification of the desired genes was carried out using ReddyMix PCR Master Mix (Thermo Scientific # AB0575DCLDB) with 120ng of genomic DNA as template using the following conditions: 95°C for 120s, 95°C for 25s, annealing (x) °C for 35 s, 72 °C for 65 s, step 2 to 4 were repeated 35 times and final extension 72 °C for 300 s where x was 53.1°C for *Rosa26*-primers, 53.6°C for NMR-specific-*Eras*-primers, and 52.4°C for *IL32*-primers. PCR amplicon was run on 1.5% agarose gel and imaged on Syngene GeneFlash Imaging system.

## Acknowledgments

FH is funded by a Gates Cambridge Trust PhD scholarship. YK is funded by a CRUK multidisciplinary award to ESS. KL is funded by a CRUK career establishment award and The Isaac Newton Trust Grant to WTK. This work was funded by generous donations from Dr. Paul Beattie, Magdalene College (Cambridge), The Isaac Newton Trust Grant (16.38c), CRUK Grant (C56829/A22053) to ESS and CRUK Grant (C47525/A17348) to WTK.

## Supplementary figures legends

**Supplementary Figure 1.**
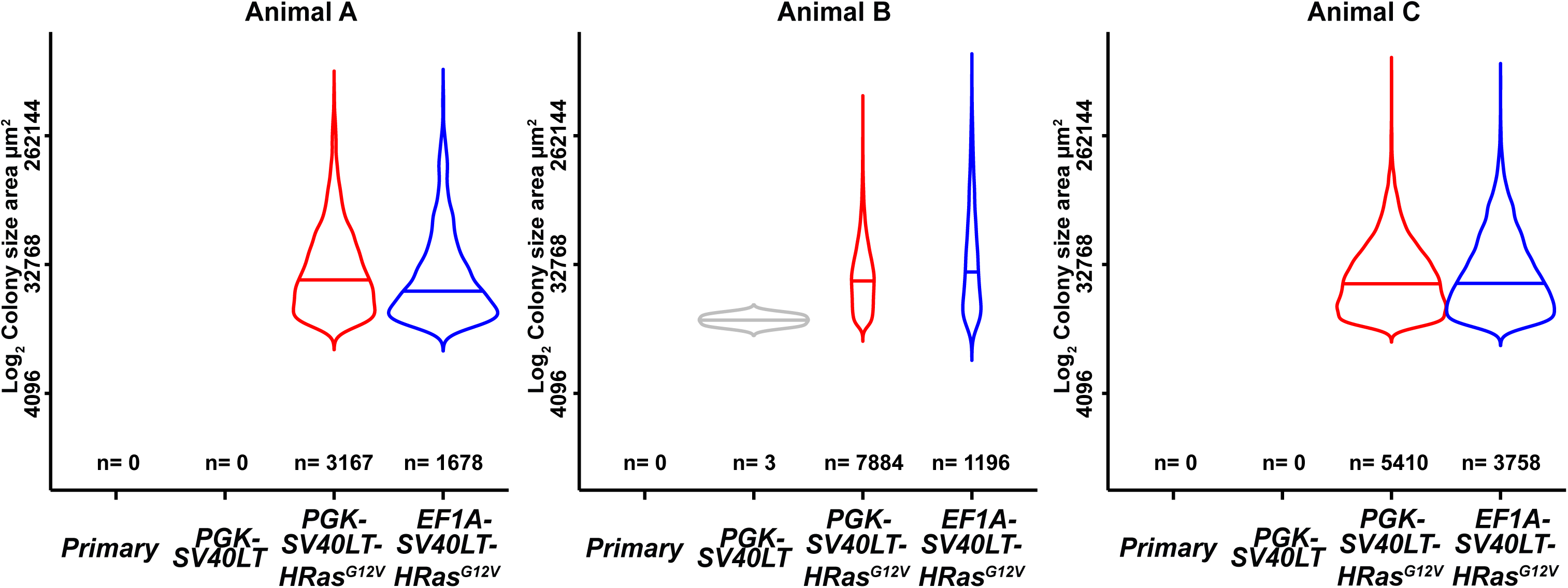
Violin plots illustrating the distribution of colony size across the different conditions for cells lines reported in Figure 1d.

**Supplementary Figure 2.**
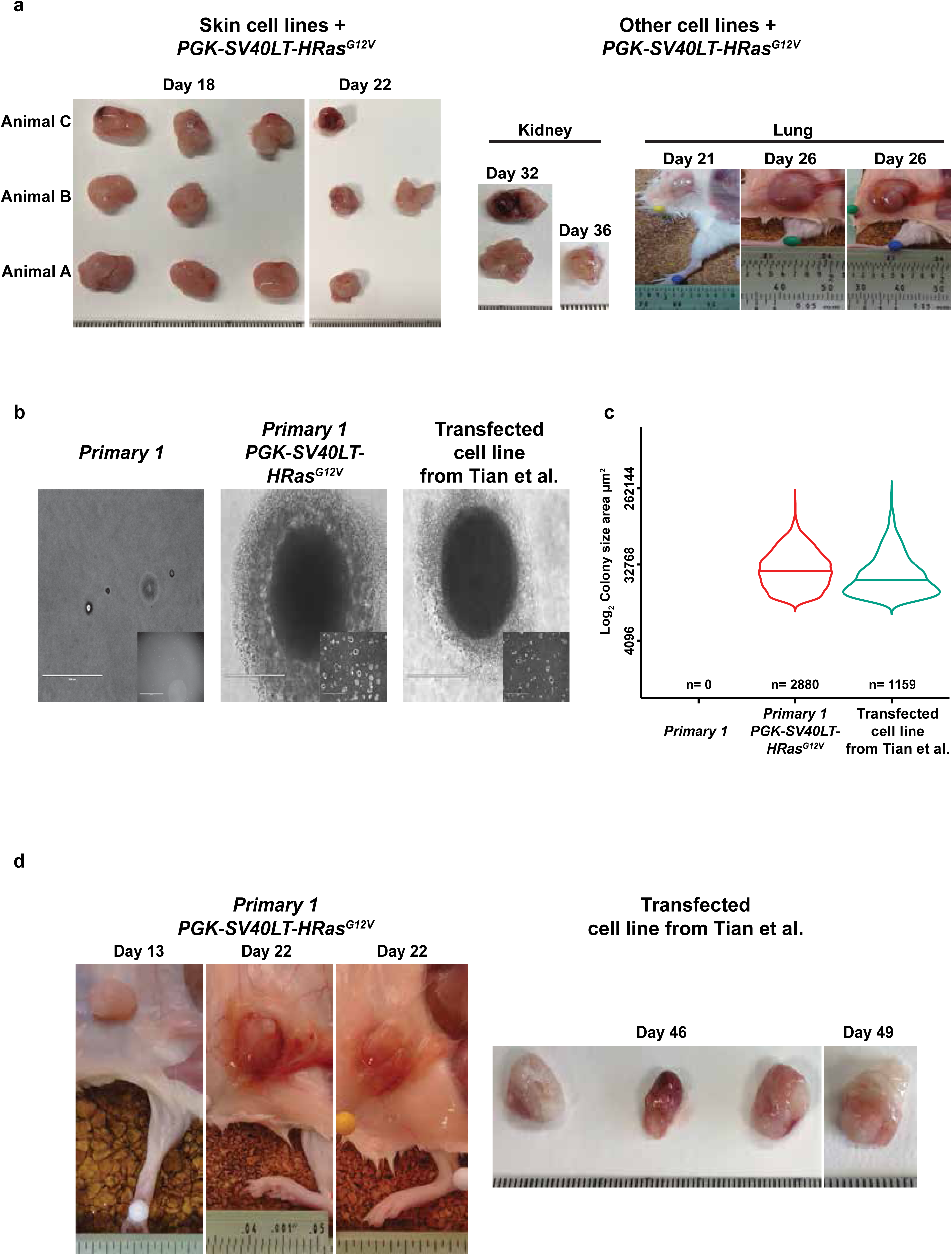
**(a)** Representative images of the xenograft tumours reported in Figure 2a-b. **(b)** Representative images of the soft agar colonies reported in Figure 2c and Figure 2e. **(c)** Violin plots illustrating the distribution of colony size across the different conditions for cells lines from Figure 2c and Figure 2e. **(d)** Representative images of the xenograft tumours reported in Figure 2d and Figure 2f.

**Supplementary Figure 3.**
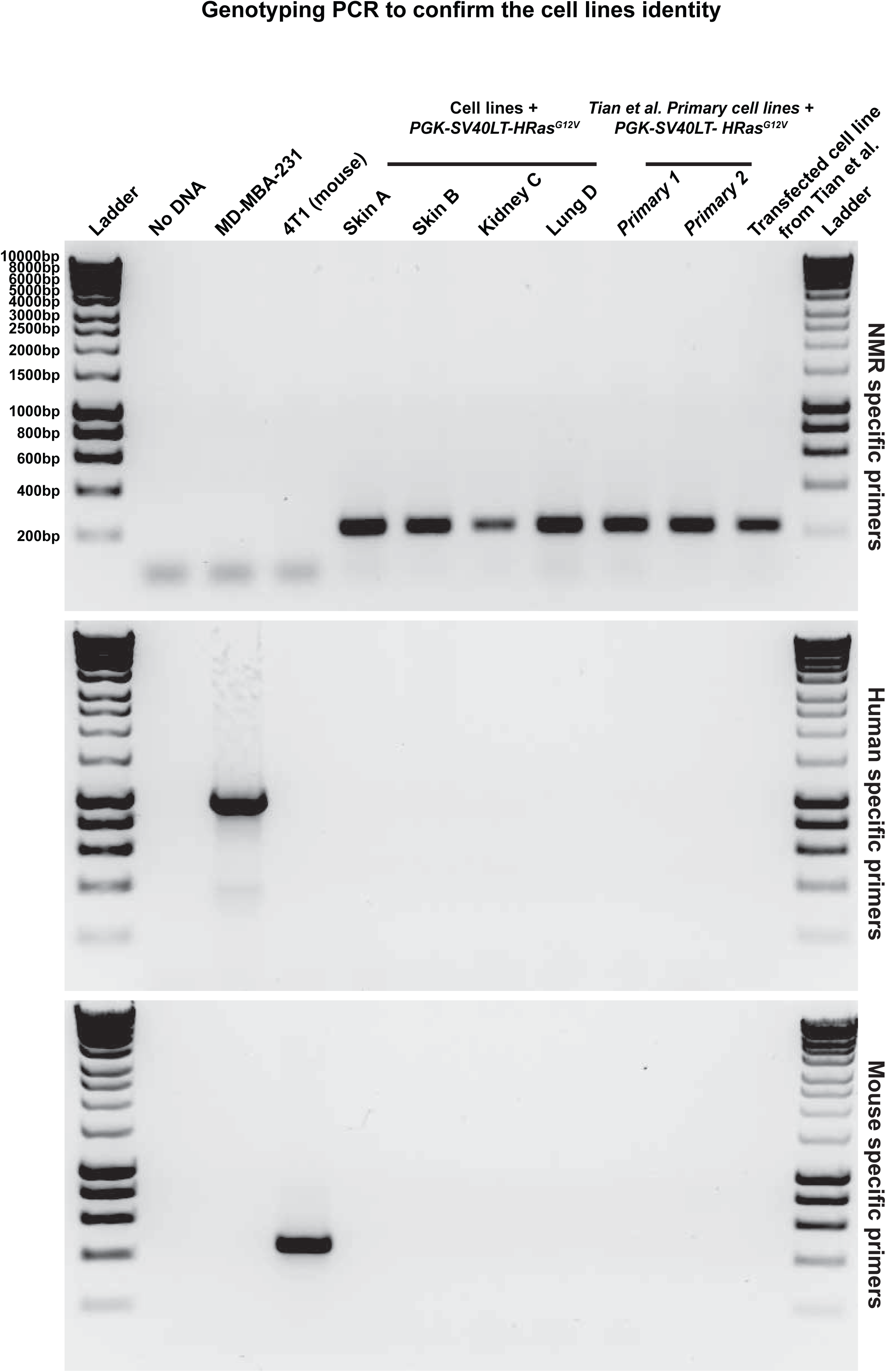
Genotyping PCR on a selection of the cell lines we generated and obtained from Tian et al. using NMR specific primers (top panel), Human specific primers (middle panel) or Mouse specific primers (bottom panel). The NMR specific primers amplify 216bp from the NMR *Eras* gene locus. The Human specific primers amplify 976bp from the Human *IL32* gene locus. The Mouse specific primers amplify 433bp from the Mouse *Rosa26* gene locus.

## Supplementary Tables

**Supplementary Table 1.** List and details of all the NMR cell lines generated. Boxes filled with green and x indicate cell line established while grey filled boxes indicate cell lines not available.

| Animal ID & Sex | Age (in weeks) at Time of Sacrifice | Weight (in grams) at Time of Sacrifice | Tissue | PGK-BFP | PGK-SV40LT | PGK-SV40LT-HRAS<sup>G12V</sup> | EF1A-SV40LT-HRAS<sup>G12V</sup> |
| --- | --- | --- | --- | --- | --- | --- | --- |
| A<br>(Female) | 244.5 | 47.5 | Skin | x | x | x | x |
|  |  |  | Pancreas |  |  |  |  |
|  |  |  | Lung |  |  |  |  |
|  |  |  | Kidney | x | X | X |  |
|  |  |  | Intestine |  |  |  |  |
| B<br>(Female) | 37.5 | 35 | Skin |  | x | x | x |
|  |  |  | Pancreas |  |  |  |  |
|  |  |  | Lung |  |  |  |  |
|  |  |  | Kidney |  |  |  |  |
|  |  |  | Intestine |  |  |  |  |
| C<br>(Male) | 58 | 60.5 | Skin | x | x | x | x |
|  |  |  | Pancreas | x | x | x | x |
|  |  |  | Lung |  |  |  |  |
|  |  |  | Kidney | x | x | x | x |
|  |  |  | Intestine |  |  |  |  |
| D<br>(Male) | 64.5 | 58 | Skin | x | x | x |  |
|  |  |  | Pancreas |  | x | x |  |
|  |  |  | Lung |  |  |  |  |
|  |  |  | Kidney |  | x | x |  |
|  |  |  | Intestine |  |  |  |  |
| E<br>(Male) | 74 | 47.5 | Skin |  |  |  |  |
|  |  |  | Pancreas | x | x | x | x |
|  |  |  | Lung | x | x | x | x |
|  |  |  | Kidney | x | x | x | x |
|  |  |  | Intestine |  |  |  |  |
| F<br>(Male) | 64 | 44 | Skin |  |  |  |  |
|  |  |  | Pancreas | x | x | x | x |
|  |  |  | Lung |  |  |  |  |
|  |  |  | Kidney | x | x | x | x |
|  |  |  | Intestine |  |  |  |  |
| G<br>(Male) | 75.5 | 43 | Skin |  |  |  |  |
|  |  |  | Pancreas | x | x | x | x |
|  |  |  | Lung | x | x | x | x |
|  |  |  | Kidney | x | x | x | x |
|  |  |  | Intestine |  |  |  |  |
| H<br>(Male) | 76.5 | 63.5 | Skin |  |  |  |  |
|  |  |  | Pancreas | x | x | x | x |
|  |  |  | Lung | x | x | x | x |
|  |  |  | Kidney | x | x | x | x |
|  |  |  | Intestine | x | x | x | x |
| I<br>(Male) | 65 | 51 | Skin |  |  |  |  |
|  |  |  | Pancreas | x | x | x | x |
|  |  |  | Lung | x | x | x | x |
|  |  |  | Kidney | x | x | x | x |
|  |  |  | Intestine | x | x | x | x |

**Supplementary Table 2.**
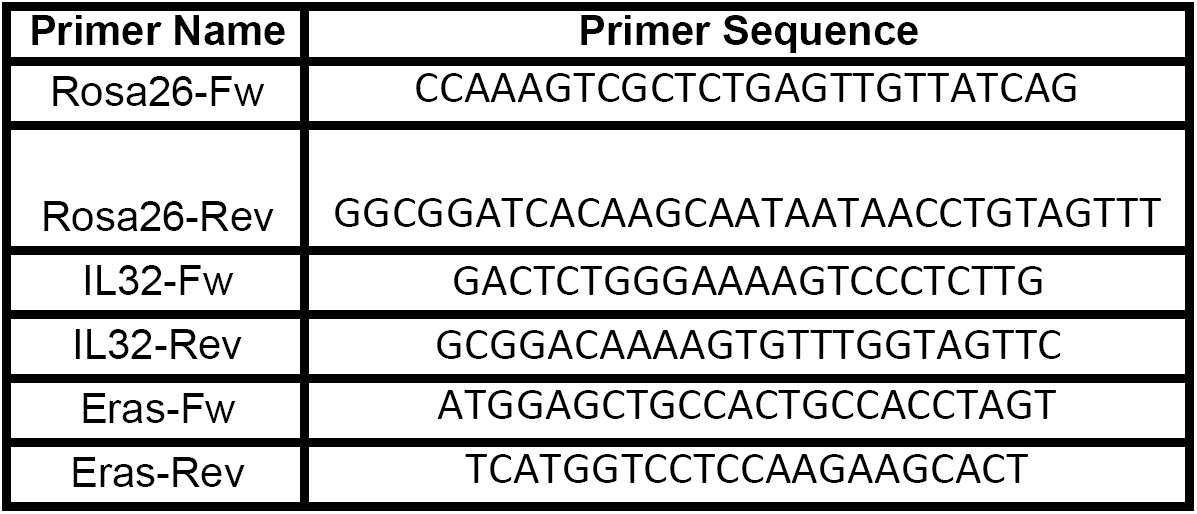
List of the genotyping PCR primers used in Supplementary Figure 3.

## References

1. Park, T. J. et al. Fructose-driven glycolysis supports anoxia resistance in the naked mole-rat. Science 356, 307–311 (2017).

2. Schuhmacher, L.-N., Husson, Z. & St John Smith, E. The naked mole-rat as an animal model in biomedical research: current perspectives. Open Access Anim. Physiol. (2015). doi:10.2147/OAAP.S5037.

3. Ruby, G. J., Smith, M. & Buffenstein, R. Naked Mole-Rat Mortality Rates Defy Gompertzian Laws and Do Not Increase With Age. Elife 7, (2018).

4. Buffenstein, R. Negligible senescence in the longest living rodent, the naked mole-rat: insights from a successfully aging species. J Comp Physiol B 178, 439–445 (2008).

5. Delaney, M. A. et al. Initial Case Reports of Cancer in Naked Mole-rats (Heterocephalus glaber). Vet. Pathol. 53, 691–696 (2016).

6. Taylor, K. R., Milone, N. A. & Rodriguez, C. E. Four Cases of Spontaneous Neoplasia in the Naked Mole-Rat (Heterocephalus glaber), A Putative Cancer-Resistant Species. J Gerontol A Biol Sci Med Sci 72, 38–43 (2017).

7. Seluanov, A. et al. Hypersensitivity to contact inhibition provides a clue to cancer resistance of naked mole-rat. PNAS 106, 19352–19357 (2009).

8. Miyawaki, S. et al. Tumour resistance in induced pluripotent stem cells derived from naked mole-rats. Nat. Commun. 7, (2016).

9. Tian, X. et al. INK4 locus of the tumor-resistant rodent, the naked mole rat, expresses a functional p15/p16 hybrid isoform. PNAS (2015). doi:10.1073/pnas.141820311.

10. Tian, X. et al. High-molecular-mass hyaluronan mediates the cancer resistance of the naked mole rat. Nature (2013). doi:10.1038/nature1223.

11. Fang, X. et al. Adaptations to a Subterranean Environment and Longevity Revealed by the Analysis of Mole Rat Genomes. Cell Rep. (2014). doi:10.1016/j.celrep.2014.07.03.

12. Kim, E. B. et al. Genome sequencing reveals insights into physiology and longevity of the naked mole rat. Nature 479, 223–7 (2011).

13. Sander, J. D. & Keith Joung, J. CRISPR-Cas systems for editing, regulating and targeting genomes. Nat. Biotechnol. 32, 3–4 (2014).

14. Rangarajan, A., Hong, S. J., Gifford, A. & Weinberg, R. A. Species-and cell type-specific requirements for cellular transformation. Cancer Cell 6, (2004).

15. Michalovitz, D., Fischer-Fantuzzi, L., Vesco, C., Pipas, J. M. & Oren’, M. Activated Ha-ras Can Cooperate with Defective Simian Virus 40 in the Transformation of Nonestablished Rat Embryo Fibroblasts. J. Virol. 61, 2648–2654 (1987).

16. Tzelepis K. et al. A CRISPR Dropout Screen Identifies Genetic Vulnerabilities and Therapeutic Targets in Acute Myeloid Leukemia. Cell Rep. 2016 Oct 18;17(4):1193–1205

